# TCF25 emerges as a cytoplasmic regulator of GPRASP2 stability

**DOI:** 10.64898/2026.04.09.714440

**Authors:** Liana J. E. Hardy, Haziq K. Moinudeen, Laura Grzegorzek, J. Fernando Bazan, Svetlana Khoronenkova

## Abstract

TCF25 is a conserved eukaryotic protein whose cellular localisation and functional context remain poorly understood. Here, we show that under basal conditions, TCF25 is predominantly cytoplasmic across several human cell types, with a digitonin-resistant perinuclear pool indicative of association to membrane-linked components. Exploration of TCF25’s proximal network using proximity labelling further supports enrichment of cytoplasmic and intracellular membrane-associated proteins. Guided by the analysis of public datasets and the proximal proteome data, we show that TCF25 associates with G-protein coupled receptor-associated sorting protein, GPRASP2, and maintains its basal abundance by protecting it from proteasomal degradation. Using structural modelling, we identify a putative interaction interface between the two proteins, involving two closely-spaced helices in the C-terminal region of TCF25 that dock to the helical repeat segment of GPRASP2. Consistent with this prediction, deletion of TCF25 amino acids 604-664 disrupted the relationship between TCF25 and GPRASP2. Together, these findings establish a cytoplasmic, membrane-proximal context for TCF25 and identify a C-terminal tract required for its association with GPRASP2. These findings implicate TCF25 in regulating GPRASP2 stability, providing a framework for investigating its role in cellular homeostasis and disease.

## Introduction

Defining the cellular localisation of a protein and how it engages specific molecular partners is central to understanding its biological function, particularly if its annotation has evolved over time [1,2]. Tcf25, also known as Nulp1, was initially described as a nuclear protein broadly expressed during mouse embryogenesis and was subsequently annotated as a putative basic helix–loop–helix (bHLH) transcription factor [3–5]. Recent functional studies of human TCF25 and its orthologues, including yeast Rqc1, have shifted this view. TCF25 is currently recognised for its conserved role in ribosome-associated quality control (RQC), where it promotes proteasomal degradation of aberrant nascent polypeptides generated during ribosome stalling [6,7]. Specifically, TCF25 facilitates Listerin-mediated K48-linked ubiquitination of stalled nascent chains on the 60S ribosomal subunit, placing it within a core proteostasis mechanism [8–10]. Other studies have linked TCF25 with nutrient-responsive signalling, lysosomal function and stress-induced cell death, indicating that it may function more broadly at the interface between proteostasis and cellular stress responses [11,12].

This functional shift has not been matched by a clear consensus on TCF25 localisation. Nuclear localisation of TCF25 and its orthologues has been mostly observed in reporter-based systems, including truncated constructs, fluorescent fusions or heterologous overexpression, and in specific biological contexts such as rat cardiomyocytes and asexual blood-stage parasites [4,12–14]. In contrast, studies examining endogenous or full-length TCF25 have reported a predominantly cytoplasmic distribution and protein associations [11,12]. Resolving this ambiguity is important because the proposed functions and emerging disease associations of TCF25 depend on the cellular compartment in which it acts. Epigenetic alterations of TCF25 have been linked to hearing loss, while high *TCF25* expression has been preliminarily associated with favourable outcomes in endometrial, glioblastoma and renal cancers [15–17].

Here, this gap was addressed using complementary cell biological, biochemical and structural modelling approaches. TCF25 was found to be predominantly cytosolic under basal conditions, and a digitonin-resistant perinuclear pool was identified. Guided by these findings and publicly available protein-association data, further investigation was focused on a predicted association between TCF25 and the G protein–coupled receptor-associated sorting protein GPRASP2. This association was confirmed and TCF25 was found to regulate GPRASP2 protein stability under basal conditions. Structural modelling and immunoprecipitation studies were then used to precisely map a C-terminal region of TCF25 that is required for this association. Together, these findings support a cytosolic function of TCF25 under basal conditions and identify a C-terminal element as essential for association with GPRASP2, thereby providing further insight into TCF25 biology.

## Results

### TCF25 localises to the cytoplasm with the membrane-proximal compartment

We initially focused on detecting the localisation of endogenous TCF25 in human cell lines of different tissue origin (Fig. 1A-C). The cell models included normal TIG1 fibroblasts, embryonic kidney HEK293-T cells, osteosarcoma U2-OS cells and human induced pluripotent stem cell (iPSC)-derived cardiomyocytes. To control for antibody specificity, TCF25 depletion using siRNA or genetic knockout (KO) was confirmed using immunoblotting and immunofluorescence (Fig. 1A-B, left and right, respectively). Across the cell lines analysed, TCF25 was predominantly distributed throughout the cytoplasm, with its signal partially overlapping with that of cytoplasmic markers including α/β-tubulin and, in differentiated cardiomyocytes, α-actinin (Figs. 1A-C and 1D, left). Selective digitonin permeabilisation of U2-OS cells, which removes soluble components and enriches for insoluble or membrane-associated components, resulted in the retention of TCF25 in the perinuclear area (Fig. 1D).

**Figure 1.**
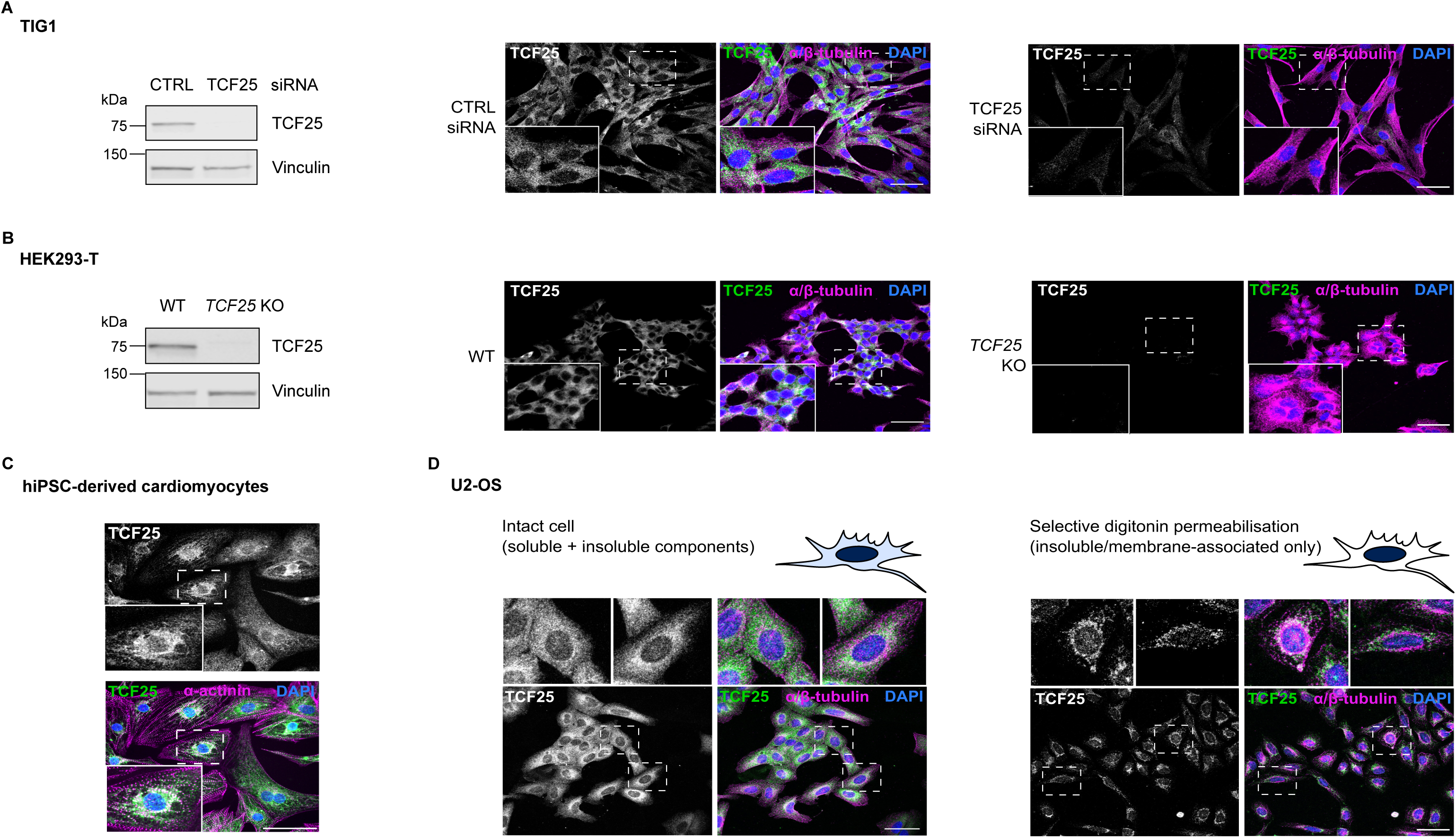
Human TCF25 is predominantly localised to the soluble cytoplasm, with a fraction in the perinuclear space. *(A, B)* Immunoblot analysis *(left)* and confocal immunofluorescence images *(right)* of *(A)* TIG1 fibroblasts treated with control (CTRL) or TCF25-targeting RNAi and *(B)* wild-type (WT) and *TCF25* knockout (KO) HEK293-T cells. Loading control: vinculin. *(C)* Pseudo-coloured single-plane immunofluorescence images of endogenous TCF25 in day 33 human induced pluripotent stem cell (hiPSC)-derived WTC mEGFP-ACTN2 cardiomyocytes expressing endogenously EGFP-tagged α-actinin. *(D)* Confocal immunofluorescence images of TCF25 localisation in U2-OS cells under basal conditions *(left)* or following permeabilisation with 0.005% digitonin *(right)*. *(A, B, D)* Compressed z-stack and *(C)* single optical plane images are shown. Green: TCF25; magenta: α/β-tubulin *(A, B, D)* or α-actinin *(C)*; blue: DAPI (DNA). Dashed boxes indicate enlarged regions. Scale bars: 50 µm. Images are representative of ≥3 independent experiments.

Furthermore, immunofluorescent imaging of endogenous TCF25 relative to mitochondrial (TOM20), endoplasmic reticulum (calnexin) and early endosomal (EEA1) markers showed partial spatial overlap. Line-scan fluorescence intensity profiles confirmed that TCF25 signal is adjacent to, but not coincident with these markers (Fig. 2A-C). Together, these observations indicate that TCF25 is predominantly a soluble cytoplasmic protein, with a perinuclear, organelle-associated distribution.

**Figure 2.**
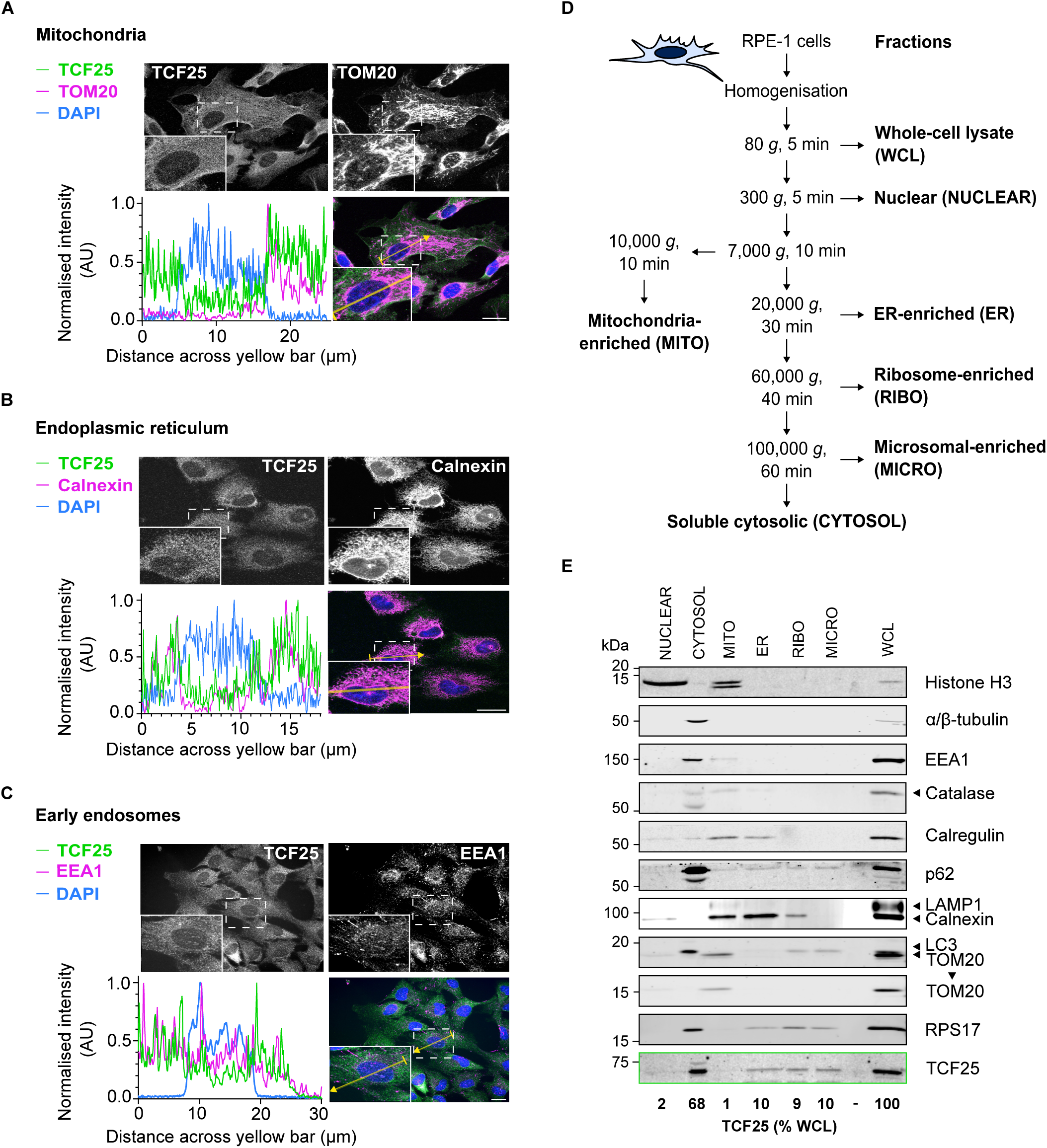
TCF25 associates with membrane-linked cytosolic compartments. *(A–C)* Immunofluorescence analysis of endogenous TCF25 localisation relative to *(A)* mitochondrial TOM20, *(B)* endoplasmic reticulum calnexin and *(C)* early endosomal EEA1 markers in RPE-1 cells. *(A, B)* Confocal single optical sections (middle slice) and *(C)* widefield single-plane images are shown. Line intensity profiles were extracted along the indicated yellow lines, background-subtracted, and normalised to each channel’s maximum to compare spatial distribution. Green: TCF25; magenta: organelle markers; blue: DAPI (DNA). Dashed boxes indicate enlarged regions. Scale bars: 10 µm. *(D)* Schematic of differential ultracentrifugation of mechanically ruptured RPE-1 cells adapted from [18,19]. TCF25 localisation was assessed in whole-cell lysate (WCL), nuclear (NUCLEAR), soluble cytosolic (CYTOSOL), mitochondria-enriched (MITO), endoplasmic reticulum-enriched (ER), ribosome-enriched (RIBO) and microsomal (MICRO) fractions. *(E)* Immunoblot analysis of TCF25 distribution across fractions as in *(D)*. TCF25 quantification is shown as a percentage of total TCF25 in WCL. Fraction enrichment using markers of chromatin (Histone H3), cytosol (α/β-tubulin), mitochondria (TOM20), peroxisomes (catalase), ER (calnexin, calregulin), endosomes/lysosomes (EEA1, LAMP1), autophagic membranes (p62/LC3) and ribosomes (RPS17) is shown.

To further define the localisation of TCF25, retinal pigment epithelial RPE-1 cells were subjected to subcellular fractionation by differential ultracentrifugation (Fig. 2D) [18,19]. This approach generated nuclear, cytosolic, mitochondria-enriched, endoplasmic reticulum (ER)-enriched, ribosome-enriched and microsomal fractions, as confirmed by the distribution of established compartment-specific marker proteins (Fig. 2D-E). Although histone H3 indicated some nuclear/chromatin carryover in the mitochondria-enriched fraction, this is consistent with the limitations of differential centrifugation and did not affect the overall pattern of TCF25 distribution [20]. Immunoblot analyses detected ∼70% of total TCF25 in the cytosolic fraction, which is consistent with the imaging data, while the remaining ∼30% enriched in ER, ribosome-enriched and microsomal-associated fractions in agreement with its roles in ribosomal and organelle biology (Fig. 2E).

Given TCF25’s distribution across soluble cytosolic and membrane-associated compartments, we next sought to define potential interaction partners of TCF25 and their subcellular localisation.

### Data mining and proximity labelling define the candidate interactome of TCF25

We initially used the Search Tool for the Retrieval of Interacting Genes/Proteins (STRING) to visualise a network of proteins with predicted and known functional and physical associations with TCF25 (Fig. S1A, left) [21]. Candidates with the highest confidence score included RQC components E3 ubiquitin ligase listerin (LTN1), an RQC cofactor NEMF and an AAA+ ATPase VCP/p97 involved in protein extraction and proteostasis [22]. Other high-score candidate associations included GPRASP2, a protein implicated in G protein-coupled receptor trafficking, and XIAP, an inhibitor of apoptosis with E3 ubiquitin ligase activity [12,23]. Noteworthy, interactions of TCF25 with the RQC components and its association with XIAP were previously confirmed using biochemical assays [8,10,12,24].

To complement this analysis, candidate TCF25-associated proteins were identified using the Human Predictome, a database of protein-protein interactions of high to moderate confidence across the human proteome, predicted and modelled by AlphaFold-Multimer [25]. Predictome-supported candidates were visualised and restricted to physical interactions using STRING, yielding a distinct set of candidate proteins (Fig. S1A, right). Of these candidate proteins, a melanoma antigen family member MAGEA11 involved in transcriptional regulation and ubiquitination and a kinase LRRK1 involved in intracellular membrane trafficking had the highest Structure Prediction and Omics Classifier (SPOC) scores. GPRASP2, however, was the only Predictome-supported candidate with a high-confidence STRING score (Fig. S1A, compare left and right). Concordant with this, all three proteins were identified as associated/interacting with TCF25 in high-throughput proteomics, yeast two-hybrid studies and other screens, with GPRASP2 reported in the highest number of independent studies curated in the Biological General Repository for Interaction Datasets (BioGRID; Fig. S1B) [26–32]. For example, in the large-scale OpenCell study, endogenously tagged TCF25 was shown to localise in the cytoplasm and GPRASP2 was detected as a stoichiometric hit in its interaction network [33].

In addition to data mining, we explored the proximal interactome of TCF25 in human RPE-1 cells using TurboID proximity labelling followed by LC–MS/MS. Biotinylated proteins proximal to TCF25 were recovered by streptavidin enrichment and quantified in TurboID-TCF25 samples relative to the whole-cell no-bait TurboID–∅ controls (Fig. S2A). The experiment was limited to two independent replicates, therefore the reproducibility threshold of Log₂ fold change > 2 in both replicates was applied, resulting in a list of 73 reproducible proximal candidates (Fig. S2B and Table S1). Among previously identified interactors of TCF25, NEMF was reproducibly enriched, however other RQC components, including LTN1 and VCP/p97, were not detected as enriched under basal conditions. In addition, GPRASP2 was enriched ∼4-fold in both TuboID replicates, although it narrowly missed the reproducibility threshold (Fig. S2B). These findings indicate that some of the previously identified interactors may be context-dependent and potentially highlight the insufficient power of our TurboID experiment. Despite these limitations, we felt that this preliminary proximal interactome study could serve as a useful exploratory dataset for other researchers in the field.

To classify reproducible proximal candidates of TCF25 according to their cellular compartments, Gene Ontology (GO) Cellular Component analysis was performed (Fig. S2C). It showed enrichment for cytoplasmic and membrane-associated compartments, such as the endomembrane system, cell junction, nuclear and ER membrane categories. Indeed, reproducible proximal candidates with the highest Log₂ fold change included the ER-resident GTPase LSG1 and the nuclear envelope protein EMD [34,35]. These findings are consistent with our imaging data and reflect the broadly perinuclear character of TCF25 proximal proteins (Figs.1D, 2A-C and S2B).

Based on the analyses of publicly available data and exploratory proximity labelling results, GPRASP2 was nominated as a proximal candidate for functional characterisation.

### TCF25 associates with GPRASP2 and influences its protein abundance

To confirm whether GPRASP2 is proximal to TCF25 under basal conditions, streptavidin pulldown fractions from TurboID experiments were probed for GPRASP2. Biotin-dependent enrichment of the bait, TurboID-TCF25, and GPRASP2 was observed in streptavidin pulldowns *vs* controls supporting their proximity in RPE-1 cells (Fig. 3A). To determine whether this proximity reflects a cellular association, co-immunoprecipitation was performed. FLAG-tagged TCF25 expressed at near-endogenous levels in U2-OS cells immunoprecipitated endogenous GPRASP2, validating an association between the two proteins (Fig. 3B).

**Figure 3.**
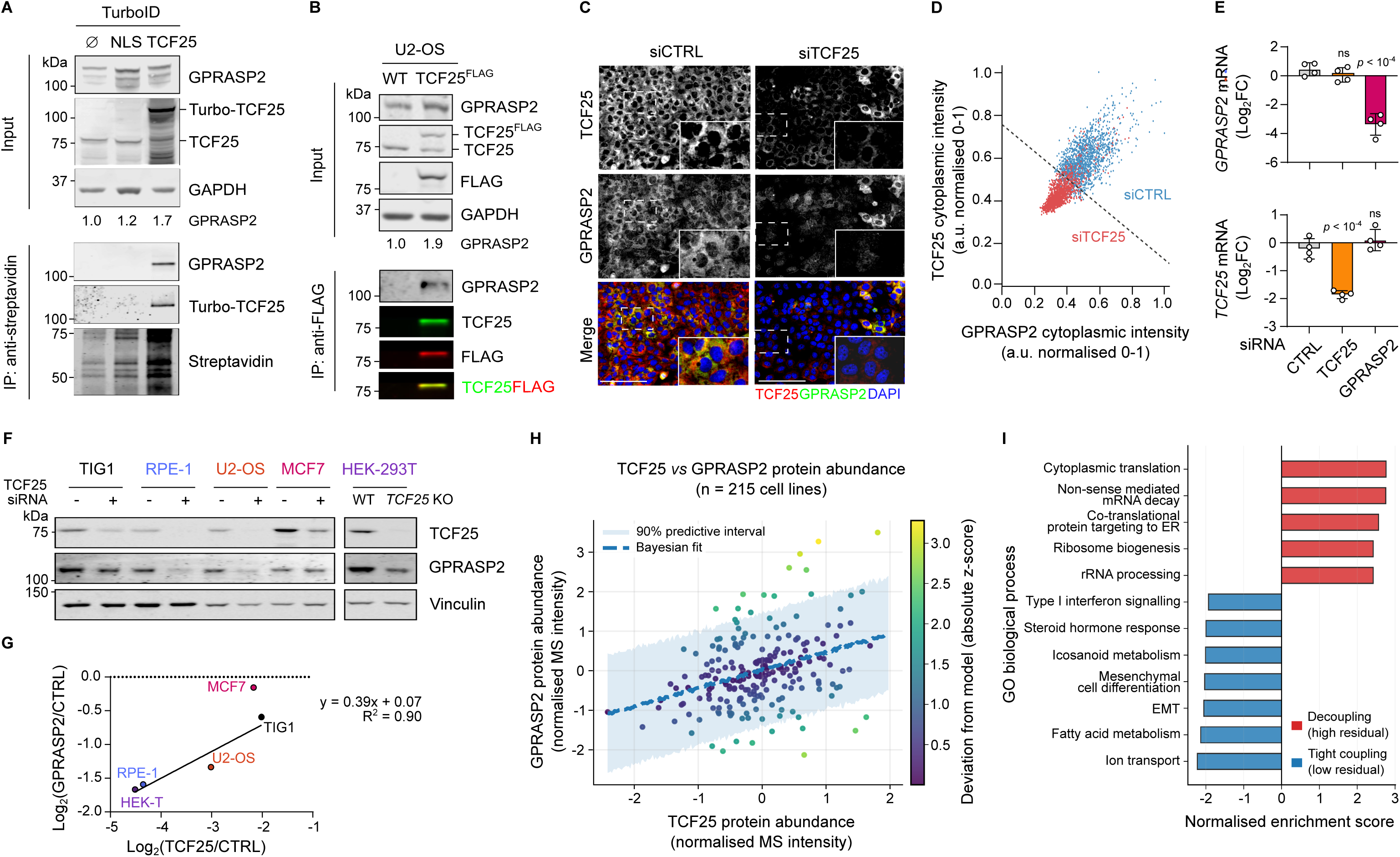
TCF25 associates with GPRASP2. *(A)* Immunoblot analysis of input and streptavidin immunoprecipitated (IP) fractions of biotinylated proteins from RPE-1 cells expressing TurboID-TCF25 or control constructs (TurboID-∅; TurboID-2xNLS). GPRASP2 levels normalised to GAPDH. Loading controls: GAPDH (input), streptavidin (IP). *(B)* Immunoblot analysis of input and anti-FLAG IPs from U2-OS cells expressing FLAG-tagged TCF25. GPRASP2 levels normalised to GAPDH. Loading control: GAPDH (input). *(C)* Immunofluorescence analysis of endogenous TCF25 and GPRASP2 protein levels following *TCF25* siRNA as in *(D)*. Red: TCF25; green: GPRASP2, blue: DAPI (DNA). Dashed boxes indicate enlarged regions. Scale bar: 20 µm. *(D)* Per-cell quantification of *(C)* displaying cytoplasmic GPRASP2 *vs* TCF25 intensity in CTRL (blue, n = 1555 cells) and TCF25 knockdown (red, n = 1478 cells) samples. Single biological replicate (10 images per condition), with each channel normalised to a 0–1 scale shown. The dashed line shows the best linear discriminant separating the two populations (accuracy = 90%). *(E)* RT–qPCR analysis of *TCF25* and *GPRASP2* mRNA levels in RPE-1 cells following non-targeting control (CTRL), *TCF25* or *GPRASP2* siRNA knockdown. Cq values normalised to *RPS13* and CTRL siRNA. Mean log₂ fold change ± SD (n = 4 independent biological experiments). One-way ANOVA with Dunnett’s multiple comparisons test. *(F)* Immunoblot analysis of TCF25 and GPRASP2 levels across several cell lines following siRNA-mediated depletion or knockout of *TCF25*. *(G)* Quantification of GPRASP2 levels relative to vinculin and TCF25 across cell lines as in *(F)*. Linear regression analysis of normalised protein abundance is shown, with the line of best fit and coefficient of determination (R²) indicated. *(H)* Bayesian correlation modelling of TCF25 *vs* GPRASP2 protein abundance across a range of established cell lines (n = 215). Quantitative protein-abundance data were obtained from the DepMap Harmonized Public Proteomics 24Q4 release of the Cancer Cell Line Encyclopedia (CCLE) proteomics dataset [36]. Each point represents one cell line and is coloured by its absolute posterior predictive z-score, indicating deviation from the fitted model. The dashed line shows the posterior mean regression; the shaded region denotes the 90% posterior predictive interval. Cell lines in *Fig. 3F-G* were missing or excluded from the analysis due to incomplete protein abundance data in the CCLE dataset. MS: mass spectrometry. *(I)* Gene Ontology (GO) Biological Process enrichment analysis of proteins with the greatest posterior predictive deviation from the Bayesian regression model. Bars represent the normalised enrichment scores (NESs). For presentation, several redundant GO terms were removed while retaining representative biological processes. ER: endoplasmic reticulum, EMT: epithelial to mesenchymal transition.

To assess whether loss of TCF25 affects GPRASP2 protein levels, single-cell level immunofluorescence analyses of cellular distributions of endogenous proteins were carried out. Levels of GPRASP2 were substantially reduced in the cells with efficient knockdown of TCF25, indicating that TCF25 may influence GPRASP2 abundance (Fig. 3C-D). To understand the mechanism of this phenomenon, we first assessed whether this effect occurs at the transcriptional level. While siRNA-mediated depletion of GPRASP2 led to a specific reduction in its transcript levels, TCF25 knockdown did not alter *GPRASP2* mRNA abundance, indicating post-transcriptional regulation of GPRASP2 levels (Fig. 3E, compare top *vs* bottom).

We further aimed to assess how widespread the correlation between TCF25 and GPRASP2 levels is across human cell lines. Reduced GPRASP2 protein levels were observed upon TCF25 loss in TIG1, RPE-1, U2-OS and HEK293-T cells (Fig. 3F). Linear regression analysis across these cell lines supported a positive relationship between TCF25 and GPRASP2 protein abundance, except in MCF7 cells (Fig. 3G). To extend this observation, quantitative mass spectrometric measurements of TCF25 and GPRASP2 protein abundance were extracted from the Cancer Cell Line Encyclopedia (CCLE) proteomics study [36]. Bayesian linear regression analysis across 215 cell lines demonstrated that TCF25 and GPRASP2 protein levels were positively correlated, with most of the cell lines falling within the 90% predictive interval (Fig. 3H). To predict biological context, in which the described positive TCF25–GPRASP2 relationship is strengthened (coupled) *vs* weakened (decoupled), posterior predictive residuals from the Bayesian model were calculated for each cell line. Cell lines were ranked by these residuals and pre-ranked GO Biological Process gene set enrichment analysis (GSEA) was performed (Fig. 3I). It was found that the decoupling of the TCF25– GPRASP2 relationship is not random and occurs preferentially in highly translationally active cell lines, where protein abundance is driven predominantly by synthesis. This finding is consistent with MCF7, a highly proliferative breast cancer line, deviating from the overall trend, although validation in a broader panel of highly proliferative lines would be required (Fig. 3G-I). In contrast, the correlation between TCF25 and GPRASP2 is most evident in stable cellular states, which are dominated by metabolic and differentiation programs (Fig. 3I).

Overall, these data indicate the existence of context-dependent post-translational regulatory mechanisms, leading us to investigate whether TCF25 may act as a scaffold or promote GPRASP2 protein stability by protecting it from proteasomal degradation.

### TCF25 regulates GPRASP2 protein stability in a proteasome-dependent manner

To investigate whether TCF25 contributes to GPRASP2 protein stability, GPRASP2 degradation was monitored after inhibition of *de novo* protein synthesis using cycloheximide (CHX) pulse-chase assays in WT and *TCF25* KO HEK293-T cells (Fig. 4A-D). Consistent with the siRNA-mediated depletion model, loss of *TCF25* resulted in significant reduction of basal GPRASP2 protein levels (Figs. 3C-D and 4A-B). Analysis of degradation kinetics showed that GPRASP2 has an estimated half-life of more than 24 h in WT cells, with measurably faster protein turnover in *TCF25* KO cells (Fig. 4C-D).

**Figure 4.**
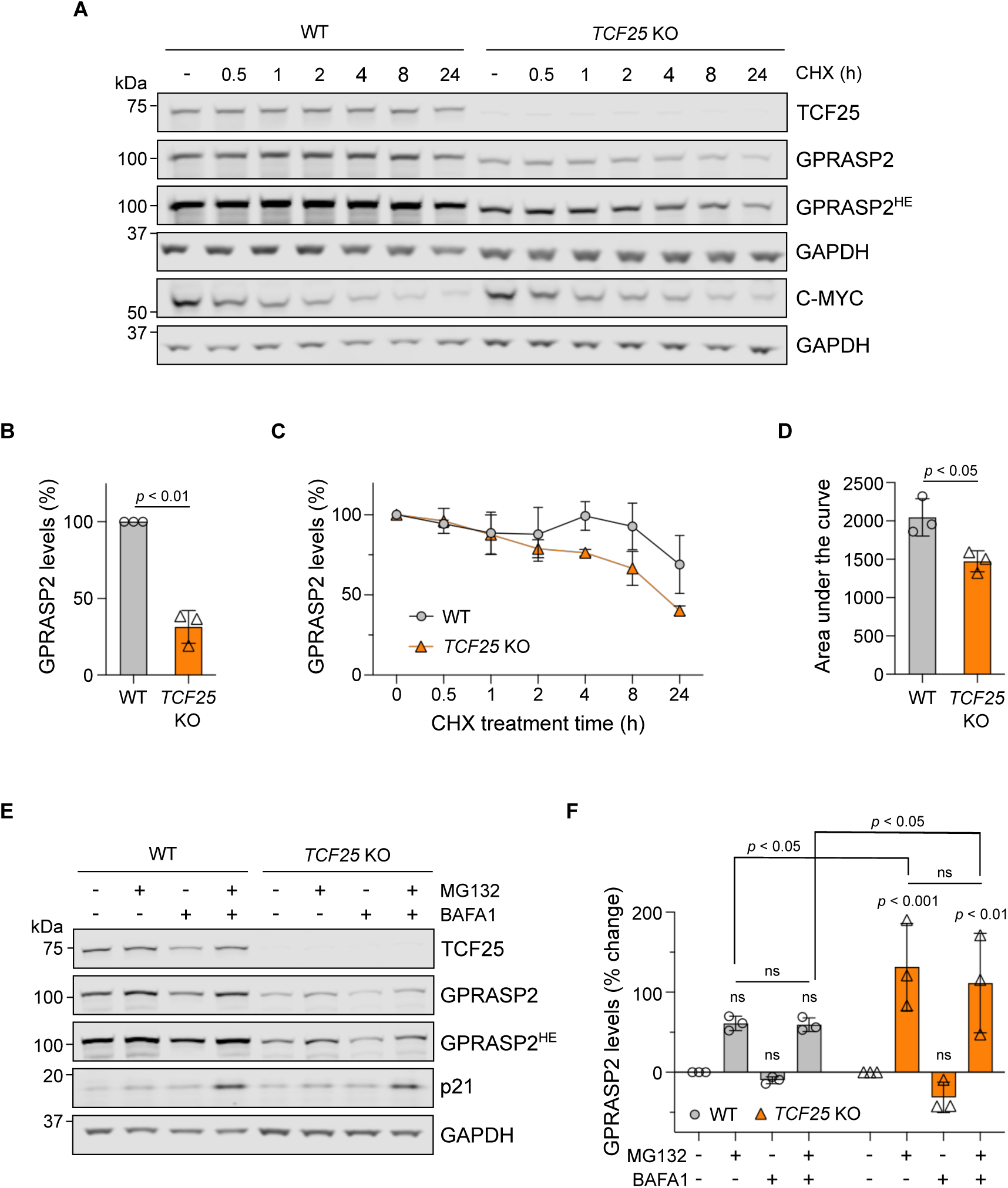
TCF25 controls basal cellular abundance of GPRASP2. *(A)* Immunoblot analysis of GPRASP2 abundance following cycloheximide (CHX) treatment of WT and *TCF25* KO HEK-293T cells. C-MYC served as a short-lived positive control (run and analysed on a separate membrane together with the loading control). Loading control: GAPDH. HE: high exposure. *(B)* Quantification of basal GPRASP2 levels in mock-treated cells as in *(A)*. Data normalised to GAPDH and WT. Mean ± SD (n = 3 independent biological experiments). One-sample *t*-test comparing to a hypothetical mean of 100. *(C)* Quantification of GPRASP2 levels as in *(A)*. Values normalised to GAPDH and respective mock-treated controls. Mean ± SD (n = 3 independent biological experiments). *(D)* Area under the curve analysis of data in *(C)*. Paired, two-tailed Student’s *t*-test. *(E)* Immunoblot analysis of GPRASP2 abundance in WT *vs TCF25* KO HEK-293T cells grown in the presence or absence of 0.5 µM MG132 and/or 100 nM bafilomycin A1 for 24 h. P21 served as a positive control. Loading control: GAPDH. HE: high exposure. *(F)* Quantification of GPRASP2 levels as in *(E)*. Data normalised to GAPDH, respective mock-treated controls and presented as % change. Mean ± SD (n = 3 independent biological experiments). Two-way ANOVA with Tukey’s multiple comparisons test.

Further, to determine the contribution of the major protein degradation pathways to GPRASP2 turnover, WT and *TCF25* KO cells were treated with the proteasomal inhibitor, MG132, the lysosomal inhibitor, bafilomycin A1 or a combination of both (Fig. 4E-F). Treatment with MG132, either alone or in combination with bafilomycin A1, increased GPRASP2 abundance in both cell lines, whereas bafilomycin A1 alone had no detectable effect. These findings indicate that basal GPRASP2 turnover is mediated predominantly by the proteasome. Notably, MG132-induced accumulation of GPRASP2 was more pronounced in *TCF25* KO than in WT cells (Fig. 4E-F).

Overall, these findings are consistent with a role of TCF25 in limiting proteasomal degradation of GPRASP2 and raise the question of which structural features mediate this association.

### The C-terminal region of TCF25 is required for its association with GPRASP2

To identify the chain segments of TCF25 that mediates its association with GPRASP2, we initially employed AlphaFold 3 (AF3)-multimer to reveal the nature and extent of the protein complex [37]. Using the full-length sequences of TCF25 and GPRASP2, AF3 narrowed the predicted interface to an approximately 80-amino-acid C-terminal segment of TCF25 that docks to the helical repeat platform of GPRASP2 (Fig. S3A). The functional importance of TCF25’s C-terminal region, which follows the major globular domain in TCF25 comprised of TPR helical repeats, is signalled by the AlphaMissense (AM) variant-effect profile. The AM profile identifies two discrete functional ‘hotspots’ that are sensitive to mutational changes, and these correspond to two α-helices (A and B) separated by a flexible linker near the TCF25 C-terminus (Fig. S3A-B). Α C-terminal 80-amino-acid peptide of TCF25 containing helices A and B is confidently predicted by AF3 to bind to grooves on the concave surface of GPRASP2, recapitulating the key features of the TCF25-GPRASP2 complex (Fig. 5A). Furthermore, AF3 defines the TCF25 peptide-GPRASP2 complex as highly confident with the predicted template modelling (pTM) and interface predicted template modelling (ipTM) scores of 0.80 and 0.84, respectively. Here, ipTM critically reflects the modelling confidence of chains at the predicted interface. These predictions were further supported by Protenix-v2, another frontier predictor, which assigned the TCF25 peptide-GPRASP2 complex high-confidence pTM and ipTM scores of 0.83 and 0.85, respectively (Fig. 5A) [38,39]. Moreover, a more recent measure of complex reliability is the interprotein surface accessible error (ipSAE), which scored the Protenix-v2 TCF25 peptide-GPRASP2 complex as highly confident with ipSAE of 0.85 [40].

**Figure 5.**
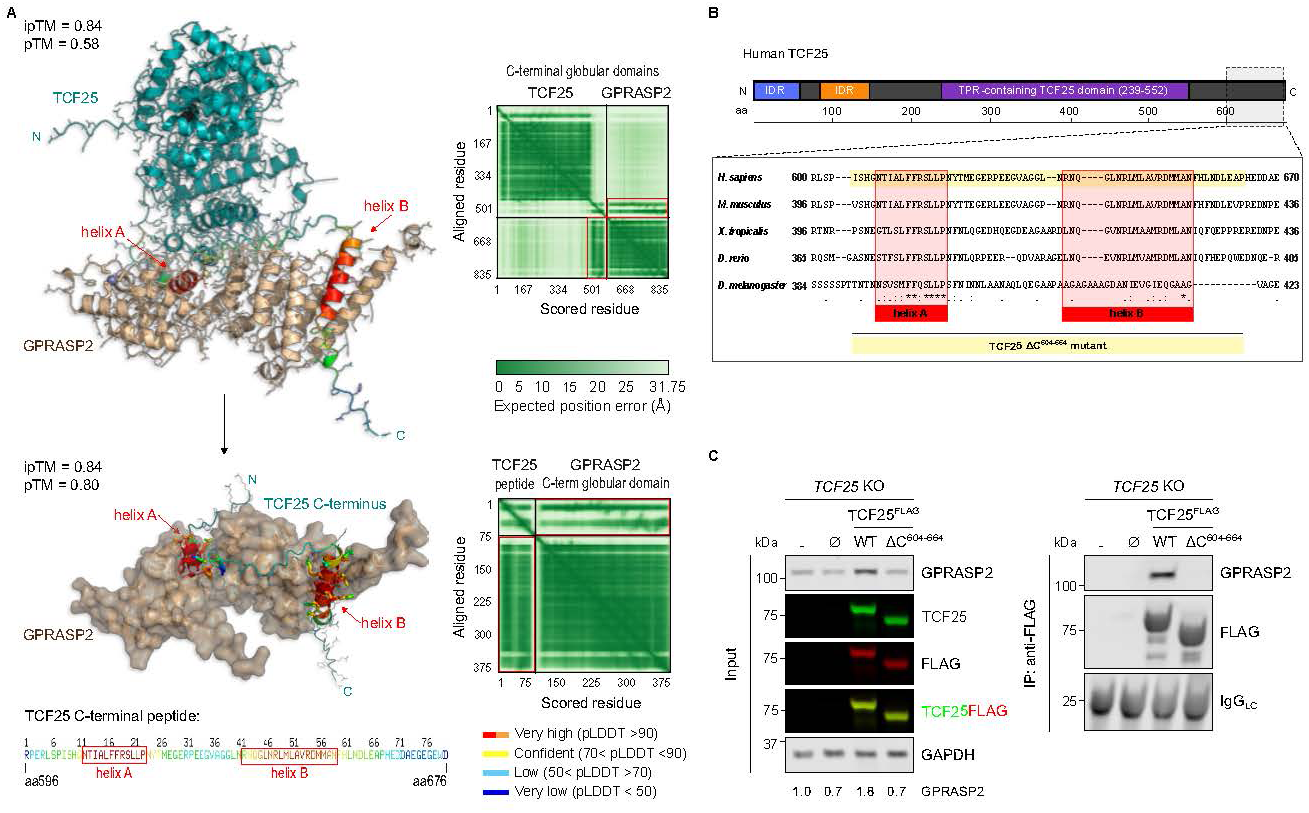
A model-guided C-terminal deletion of TCF25 disrupts interaction with GPRASP2. *(A)* Predicted C-terminal globular domains of TCF25 and GPRASP2 as in *Fig. S3* were confidently predicted to form a heterodimeric complex *(top left)*. The second model shows the TCF25 C-tail peptide (amino acids 596-676), narrowing the predicted interaction elements with GPRASP2 *(bottom left)*. Predicted template modelling (pTM) and interface predicted TM (ipTM) scores *(left)* and predicted aligned error (PAE) plots for both models are shown *(right)*. The TCF25 C-tail peptide is coloured by Predicted Local Distance Difference Test (pLDDT) scores, where red-orange-yellow paint highlights residues that are confidently predicted at interfaces, with helices A and B highlighted in red boxes. Models generated using the AlphaFold v3.2 [37] and Protenix-v2 servers [38]. *(B)* Multiple sequence alignment of vertebrate TCF25 C-terminal protein sequences. Degree of conservation is annotated: fully conserved residues (*), strongly conserved substitutions (:) and weakly conserved substitutions (.). *(C)* Immunoblot analysis of input and anti-FLAG IPs from HEK-293T *TCF25* KO cells expressing FLAG-tagged WT or the C-terminal deletion mutant (ΔC^604-664^) of TCF25. Untransfected cells (**–**) and cells expressing empty plasmid construct (∅) served as controls. GPRASP2 levels normalised to GAPDH. Loading controls: GAPDH (input), immunoglobulin light-chain, IgGLC (IP).

The evolutionary conservation of the predicted interface is highlighted by a cross-species alignment of the TCF25 C-terminal segment (Fig. 5B). The analysis identified conserved residues clustering in the two recognition α-helices across several species, consistent with their predicted functional role. Based on the structural predictions and sequence analysis, a TCF25 deletion mutant lacking amino acids 604-664 and encompassing both predicted α-helices, was prioritised for experimental assessment of the predicted TCF25-GPRASP2 interface (Fig. 5B).

To this end, FLAG-tagged WT TCF25 or the C-terminal deletion mutant ΔC^604-664^ was expressed in *TCF25* KO HEK-293T cells, followed by FLAG pull-down assays (Fig. 5C). GPRASP2 protein levels were modestly increased following WT TCF25 expression, consistent with observations in RPE-1 cells expressing TurboID-TCF25 and U2-OS cells expressing WT TCF25 (Figs. 3A-B and 5C). The ΔC^604-664^ mutant was expressed at a level comparable to that of WT TCF25, albeit it did not increase GPRASP2 abundance. In agreement with this, FLAG-tagged WT TCF25 co-immunoprecipitated with endogenous GPRASP2, whereas no GPRASP2 was detected in ΔC^604-664^ pull-downs (Fig. 5C). These data are consistent with the structural predictions and support a requirement for the TCF25 C-terminal region for its association with GPRASP2.

Together, these findings support a functional association between TCF25 and GPRASP2 and identify TCF25 amino acids 604-664 as important for GPRASP2 association and stabilisation.

## Discussion

Previous reports have described variable localisation of TCF25 across experimental systems, including nuclear and cytoplasmic, as well as context-dependent relocalisation to cytoplasmic organelles such as ribosomes and lysosomes [3,4,9,11,12]. Our data resolve this ambiguity by demonstrating that, under basal conditions, TCF25 is largely cytoplasmic and diffusible (Figs. 1 and 2). We also identify a smaller, membrane-proximal fraction associated with perinuclear organelle compartments. This distribution is similar across several human cell lines and supported by imaging with selective digitonin-based permeabilisation. Consistent with this, biochemical fractionation identifies predominantly cytoplasmic localisation of TCF25, with a smaller pool in ER-, ribosome- and microsome-enriched fractions (Fig. 2). These observations agree with the general cytoplasmic residence reported in recent analyses of endogenous TCF25 localisation [11], but differ from initial sequence- and overexpression-based classification of TCF25 as a transcription factor [4,13]. Contemporary sequence and domain analyses do not support the presence of a canonical DNA-binding bHLH module [5,41]. Instead, TCF25 appears to contain a basic, intrinsically disordered N-terminus and a structured C-terminal tetratricopeptide repeat (TPR)-like helical solenoid, an architecture more consistent with protein–protein interface assembly than sequence-specific DNA binding [41,42]. Systematic classifications of human transcription factors similarly do not support TCF25 as a likely sequence-specific transcription factor [43]. Instead, reports of transcriptional changes following TCF25 perturbation may reflect indirect downstream effects rather than direct transcriptional regulation [13,14]. Despite this, regulated or transient cell-, isoform- and context-specific access of TCF25 to the nucleus cannot be excluded.

The cytoplasmic and ribosome fraction-enriched distribution of TCF25 together with the detection of NEMF in its exploratory proximal interactome are consistent with its established roles in RQC [8,9,24]. The absence of other RQC components may reflect transient or stress-dependent nature of RQC complex assembly, although the limited replication of the TurboID experiment should also be considered. This localisation however does not exclude alternative, RQC-independent functions.

Within this cytoplasmic, membrane-proximal context, our data highlight a functional association between TCF25 and GPRASP2 (Fig. 3). GPRASP2 is an X-chromosome-encoded protein and GPRASP family proteins have been implicated in post-endocytic sorting of internalised G protein–coupled receptors, facilitating their targeting to degradative compartments and contributing to receptor turnover and signal attenuation [23,44,45]. Loss of TCF25 reduced GPRASP2 protein abundance, without affecting the levels of its mRNA, and accelerated protein turnover. In addition, proteasome inhibition led to stabilisation of GPRASP2 preferentially in TCF25-deficient cells, supporting a role for TCF25 in limiting its proteasomal degradation. Although the exact mechanism of this effect requires further investigation, TCF25 may stabilise GPRASP2 by maintaining it within a protein complex or restricting access by an E3 ubiquitin ligase [46]. This stabilising activity appears distinct from the role of TCF25 in promoting degradation of aberrant nascent chains during RQC [6,8,9,24]. Overall, it is possible that TCF25 may more broadly influence the turnover of client proteins associated with intracellular membranes beyond GPRASP2 [47,48]. This hypothetical function would be consistent with reports implicating TCF25 in adaptive proteostatic responses to nutrient stress, where it enhances lysosomal acidification to support metabolic adaptation [10].

Deep learning structure prediction and deletion analysis identified TCF25 amino acids 604– 664 as necessary for its association with GPRASP2. Deletion of this semi-conserved C-terminal region, which contains two predicted recognition α-helices, abolished detectable GPRASP2 co-immunoprecipitation and stabilisation. These findings establish a requirement for the C-terminal region of TCF25 in mediating the association with GPRASP2, although the direct interaction of the proteins requires further evidence. The functional importance of this region is supported by previous analyses. C-terminal truncations of human TCF25 impaired its ability to impose K48-linkage specificity during LTN1-mediated ubiquitination, with amino acid residues 638-642 important for the interaction with acceptor ubiquitin [8]. One possibility is that TCF25 engages distinct partners in different cellular pools, including ribosome-associated complexes *vs* stabilising GPRASP2 in a membrane-proximal context. Alternatively, deletion of residues 604-664 may disrupt a shared structural module rather than a GPRASP2-specific interface. In addition, disruption of the *D. melanogaster* orthologue Nulp1 extended C-terminal region caused partial lethality during the development and bent femurs in surviving flies, with altered Wingless/Wnt-target gene expression [49]. Together, these findings link TCF25 structural features to its ability to stabilise GPRASP2, providing a mechanistic basis for this association.

In summary, we propose that TCF25 functions as a predominantly cytoplasmic, membrane-proximal scaffold that stabilises GPRASP2 and this function may influence GPCR-dependent context-specific signalling [15–17]. More broadly, TCF25 may act as a regulator of protein stability, and defining the consequences of its loss of function will be key to understanding novel TCF25-mediated signalling pathways in cellular homeostasis and disease.

## Experimental procedures

### Reagents and resources

All reagents, cell lines, and software used in this study are listed in Table 1 (Resource table).

**Table 1.**
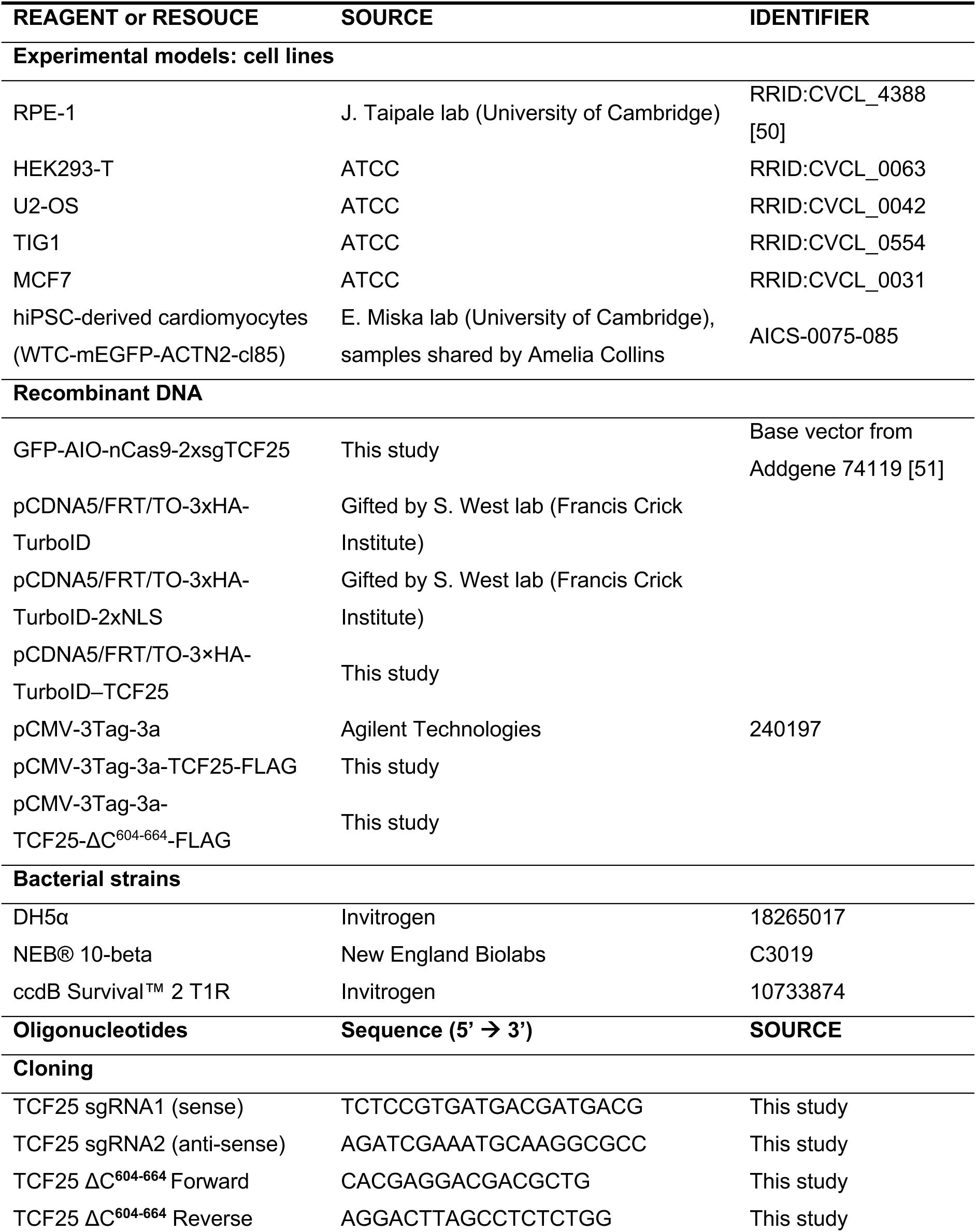
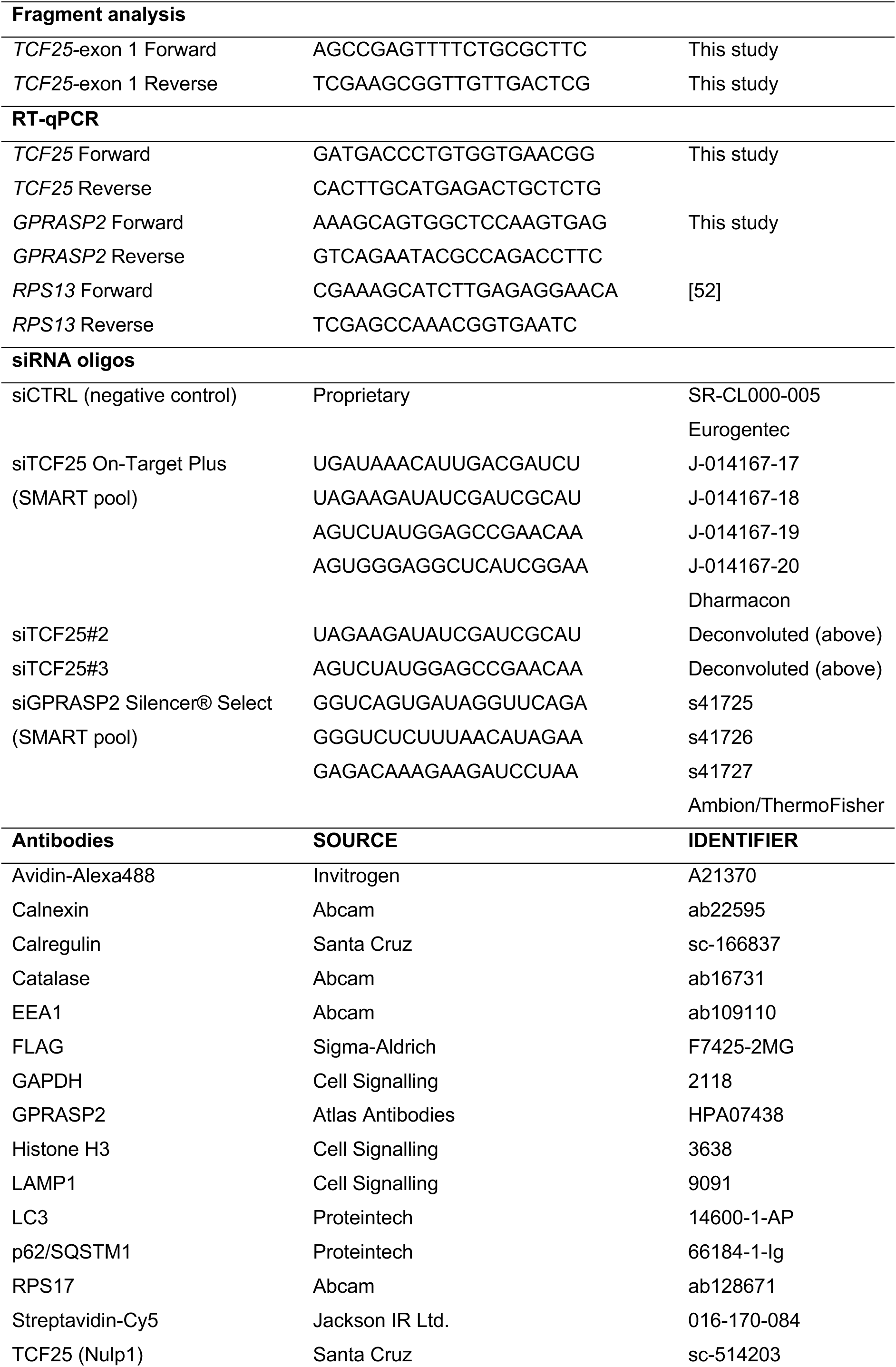
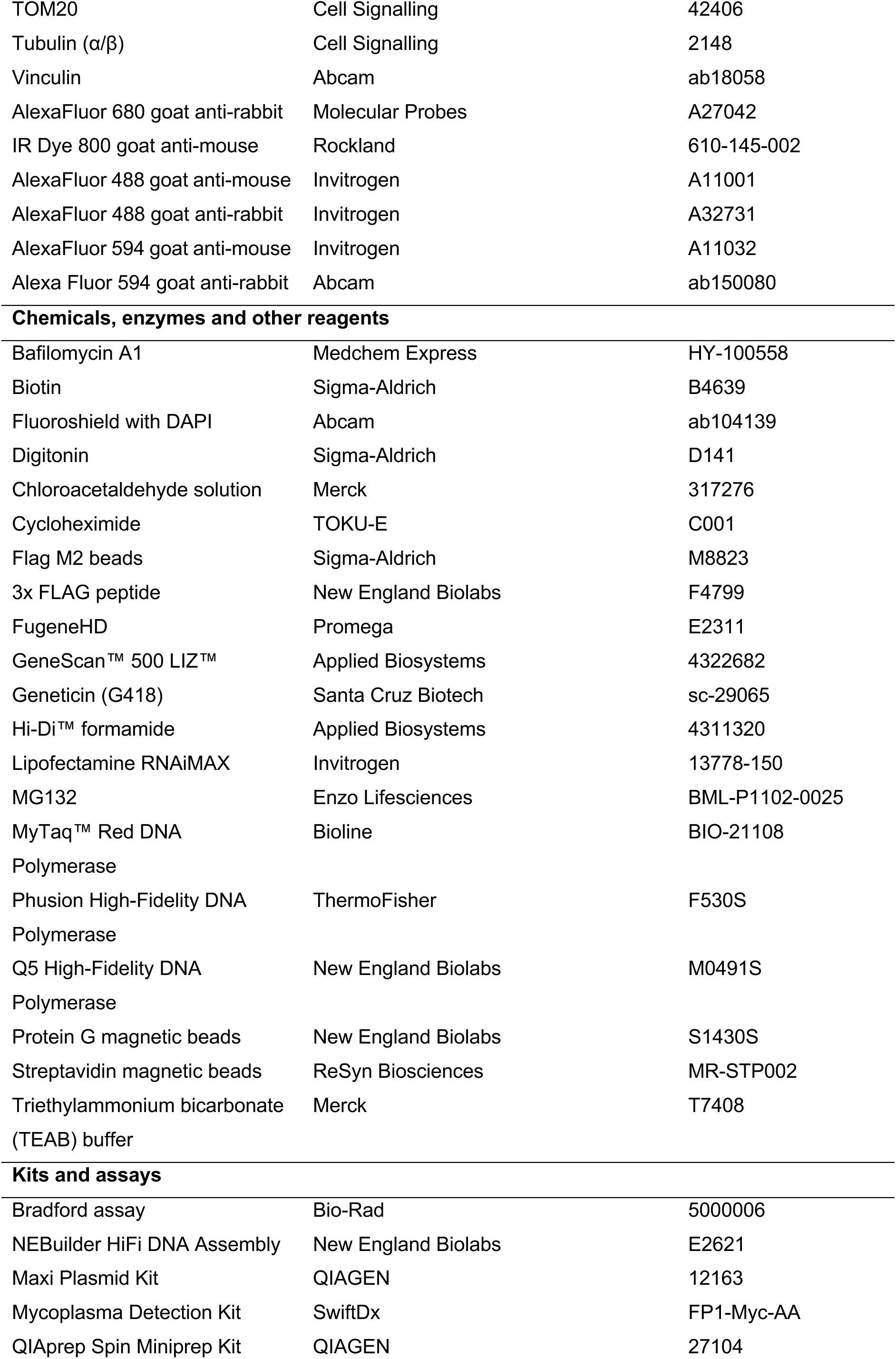
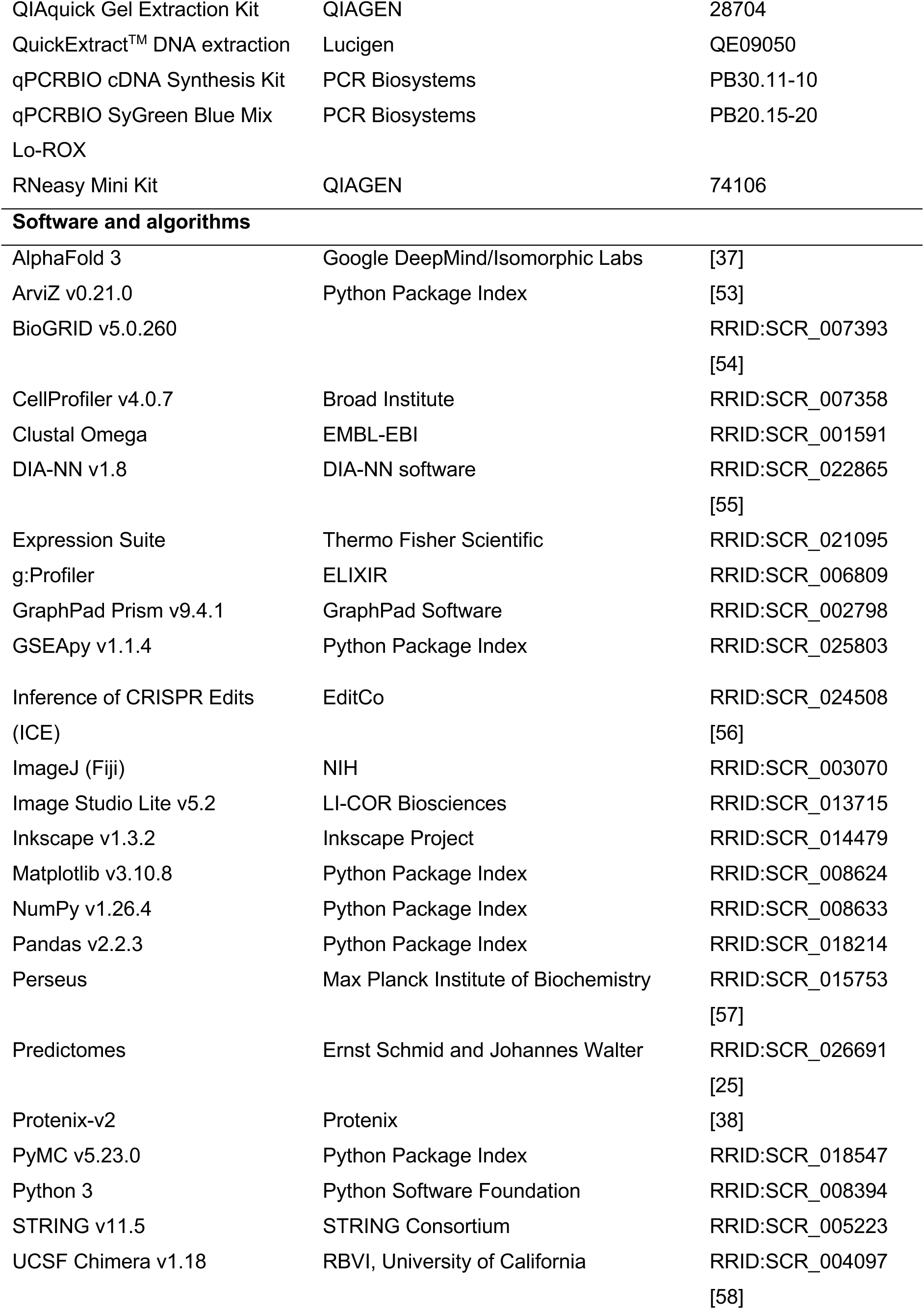
Resource table.

### Cell culture

Human hTERT-immortalised retinal pigment epithelial RPE-1 cells were obtained from the Taipale laboratory and maintained in DMEM/F-12 (Gibco™) supplemented with 10% (v/v) foetal bovine serum (FBS; Gibco™). HEK293-T cells and derivative lines were cultured in DMEM (4.5 g/L glucose, no pyruvate; Gibco™) supplemented with 10% (v/v) FBS. U2-OS osteosarcoma cells were cultured in DMEM (low glucose; Gibco™) supplemented with 10% (v/v) FBS and sodium pyruvate. U2-OS FLAG–TCF25 stable cell lines were maintained in the same medium supplemented with 0.2 mg/mL G418. TIG1 human fibroblasts were cultured in DMEM supplemented with 15% (v/v) FBS. MCF7 breast cancer cells were cultured in DMEM supplemented with 10% (v/v) FBS. Human induced pluripotent stem cell (hiPSC)-derived cardiomyocytes (WTC mEGFP-ACTN2 line, Allen Institute for Cell Science) were maintained by Amelia Collins (Miska lab, University of Cambridge) as previously described and according to supplier recommended conditions [59].

All cell lines were cultured at 37 °C in 5% CO₂ and 95% humidity. Cells were routinely tested for mycoplasma contamination and experiments were performed below passage 25–35 depending on cell line.

### Generation of TCF25 KO HEK293-T cells

*TCF25* knockout (KO) HEK293-T cells were generated using a CRISPR/Cas9 nickase (D10A)-based strategy as described previously [51]. Paired single guide RNAs (sgRNAs) targeting exon 1 of TCF25 were delivered using an all-in-one plasmid encoding Cas9 and a GFP reporter. HEK293-T cells were transfected using polyethylenimine (PEI) at a 3:1 of PEI:DNA ratio, and GFP-positive cells were enriched by fluorescence-activated cell sorting (FACS) followed by single cell expansion.

Monoclonal cell lines were screened by fluorescent PCR–capillary electrophoresis adapted from a published method [60]. Genomic DNA was extracted using QuickExtract™ DNA Extraction Solution (Lucigen), and a region flanking the sgRNA target site was amplified using fluorescently labelled forward primers (6-FAM for parental samples and HEX for edited samples). PCR was performed using Phusion High-Fidelity DNA Polymerase (ThermoFisher) with annealing at 68 °C for 35 cycles, with all other conditions according to the manufacturer’s protocol. Amplicons were combined with Hi-Di formamide and GeneScan 500 LIZ size standard and resolved using a 3100 XL Genetic Analyser (Life Technologies) at the Department of Biochemistry sequencing facility (University of Cambridge, UK).

Successful knockout was confirmed by Sanger sequencing and indel analysis using ICE software, and loss of TCF25 protein expression was validated by immunoblotting [56]. A validated monoclonal cell line (#E9), harbouring a frameshift-inducing indel in exon 1 predicted to generate a truncated, non-functional protein, was used for experiments.

### Generation of U2-OS TCF25–FLAG cell lines

U2-OS cells stably expressing TCF25-FLAG were generated by transfection with pCMV-3Tag-3a-TCF25-FLAG followed by selection with geneticin (200 µg/mL) to isolate cells with random genomic integration of the expression construct. Resistant cells were expanded and monoclonal cell lines were isolated. Clones were screened for TCF25-FLAG expression by immunoblotting and immunofluorescence, and a clone expressing TCF25-FLAG at near-endogenous levels was selected for subsequent experiments.

### RNA interference

Small interfering RNA (siRNA) duplexes were synthesised by Merck and used at a final concentration of 10 nM unless otherwise stated. Deconvoluted TCF25 siRNA duplexes (siTCF25#2 and siTCF25#3) were used at a combined final concentration of 10 nM. siRNA was delivered by reverse transfection using Lipofectamine RNAiMAX (Invitrogen) according to the manufacturer’s instructions. Culture medium was replaced after 24 h, and downstream analyses were performed 72 h post-transfection, when cell density reached ∼70–80% confluence.

### Chemical treatments

HEK293-T cell lines were treated with the proteasomal inhibitor MG132 (Enzo Life Sciences) at 0.5 µM or the lysosomal inhibitor bafilomycin A1 (MedChemExpress) at 100 nM for 24 h. For protein half-life analyses, cells were treated with 50 μg/mL cycloheximide (CHX) for varying durations up to 24 h. Control cells were treated with an equivalent volume of DMSO, at a final concentration of <0.1% (v/v). All treatments were performed under standard culture conditions.

### RT-qPCR

RT-qPCR analysis was performed as previously described [61]. Briefly, total RNA was extracted using the RNeasy Mini Kit (QIAGEN) and treated with DNase I (Thermo Fisher Scientific) according to the manufacturer’s instructions. cDNA was synthesised from 400 ng intact RNA using the qPCRBIO cDNA Synthesis Kit (PCR Biosystems). Quantitative PCR was performed using qPCRBIO SyGreen Blue Mix Lo-ROX (PCR Biosystems) on a QuantStudio™5 Real-Time PCR System (Thermo Fisher Scientific) using gene-specific primers (Table 1). Relative gene expression was calculated using the ΔΔCt method with *RPS13* as a reference gene, log₂-transformed and expressed as fold change relative to the appropriate control.

### Immunofluorescence and pre-extraction

Cells grown on glass coverslips were fixed in 4% paraformaldehyde (PFA) followed by permeabilisation with 0.2% (v/v) Triton X-100 in PBS for 10 min or alternatively fixed and permeabilised simultaneously using 100% ice-cold methanol. Coverslips were blocked in 3% (w/v) bovine serum albumin (BSA), 0.05% (v/v) Tween®20, and 0.3 M glycine in PBS for 1 h. Samples were incubated with primary antibodies overnight at 4 °C, washed in 0.05% (v/v) Tween®20 in PBS and incubated with fluorophore-conjugated secondary antibodies for 45 min at room temperature. Coverslips were washed as above and mounted onto glass slides using DAPI-containing mounting medium (Abcam).

For selective removal of soluble cytosolic material, coverslips were incubated for 5 min in digitonin-containing extraction buffer (20 mM HEPES, pH 7.6, 110 mM potassium acetate, 2 mM magnesium acetate, 0.5 mM EGTA, 2 mM DTT, 1 µg/mL leupeptin, pepstatin and aprotinin, 0.005% digitonin) prior to fixation in 100% ice-cold methanol, enriching insoluble or membrane-associated cellular structures.

### Microscopy and image analysis

Fluorescence imaging was performed using Zeiss Axio Imager M2 or Axio Imager Z2 epifluorescence microscopes equipped with 20× or 40× objectives. Confocal imaging was performed using a Nikon Eclipse Ti microscope with 40× or 60× oil immersion objectives. Widefield images were acquired as single optical sections, whereas confocal images were acquired as z-stacks and displayed as either single central sections or maximum intensity projections as indicated. Acquisition settings were kept constant within experiments.

Line scan intensity profiles were generated from single-plane images using Fiji (ImageJ). Line regions of interest (ROIs; 5-pixel width) were drawn across representative cells and applied identically to each channel. Fluorescence intensity values were extracted for each channel, and background subtraction was performed by subtracting the minimum intensity value within each channel profile. Intensity profiles were then normalised to the maximum value for each channel (min–max normalisation, 0–1) and plotted as a function of distance to assess relative spatial distribution and overlap of signals. Representative images are shown in the figures.

For per-cell cytoplasmic intensity, nuclei were segmented using DAPI staining in CellProfiler, and cell boundaries were approximated by a fixed 10-pixel isotropic expansion from each nucleus. Subtracting the nuclear mask from this region defined a perinuclear cytoplasm compartment used to quantify GPRASP2 and TCF25 signal. Mean cytoplasmic intensity was measured per cell for each channel from a single biological replicate, with 10 images acquired per condition (siCTRL: 1555 cells; siTCF25: 1478 cells). For comparison of siCTRL and siTCF25 knockdown, per-cell GPRASP2 and TCF25 intensities were each independently rescaled to a 0–1 range by dividing by the maximum single-cell value for that channel within the plotted dataset.

### Preparation of whole-cell lysates

Cells were lysed on ice in RIPA buffer (25 mM Tris–HCl, pH 8.0, 150 mM NaCl, 1% (v/v) Triton X-100, 1% (w/v) sodium deoxycholate, 0.1% (w/v) sodium dodecyl sulphate (SDS), 5 mM EDTA) supplemented with 1 mM phenylmethylsulfonyl fluoride, 1 μM staurosporine, 1 μg/ml each of aprotinin, chymostatin, leupeptin, pepstatin, *N*-ethylmaleimide, 1 µM MG132 and 1× phosphatase inhibitor cocktail (Calbiochem). Lysates were incubated for 30 min at 4 °C with rotation and centrifuged at 20,000 *g* for 20 min at 4 °C. The protein concentration of supernatants was measured using Bradford assay reagent (Bio-Rad). Lysates were prepared at equal concentrations and boiled at 95 °C for 10 min in denaturing loading buffer (25 mM Tris–HCl, pH 6.8, 2.5% v/v β-mercaptoethanol, 1% (w/v) SDS, 5% (v/v) glycerol, 0.05 mg/ml bromophenol blue, 1 mM EDTA) prior to SDS-PAGE.

### Subcellular fractionations

Subcellular fractionation was performed by differential ultracentrifugation adapted from established mitochondrial and organelle isolation workflows [18,19]. All procedures were carried out at 4 °C using Ca²⁺- and Mg²⁺-free buffers to preserve organelle integrity.

Cells were harvested by trypsinisation, washed in ice-cold PBS, and resuspended in hypotonic sucrose buffer (30 mM Tris–HCl, pH 7.4, 225 mM mannitol, 75 mM sucrose, 0.1 mM EGTA) supplemented with inhibitors (as above). Cells were lysed using a ball-bearing homogeniser (12 µm clearance; Isobiotec) with ∼20 passes to disrupt the plasma membrane while maintaining intracellular organelles intact.

Unlysed cells and debris were removed by low-speed centrifugation (80 *g*, 5 min, repeated twice), followed by centrifugation at 300 *g* for 5 min to obtain the nuclear fraction. The post-nuclear supernatant was subjected to sequential centrifugation at increasing speeds to enrich subcellular fractions: crude mitochondrial/heavy membrane fractions (7,000–10,000 *g*), intermediate membrane fractions (20,000–60,000 *g*), and microsomal fractions (100,000 *g*). The final supernatant was collected as the cytosolic fraction. The protein content of cytosolic fractions was determined by Bradford assay, and equal volumes of all fractions were analysed by SDS–PAGE, maintaining a 1:1 loading ratio between fractions.

### SDS–PAGE and immunoblotting

Proteins were separated using 4–16% Tris–glycine SDS–PAGE gels and transferred onto Immobilon®-FL PVDF membranes (Merck). Membranes were blocked using Intercept®-TBS blocking buffer (LI-COR Biosciences). Primary antibodies were diluted in Intercept®-TBS containing 0.1% (v/v) Tween®20, and secondary antibodies were diluted in the primary antibody dilution buffer supplemented with 0.01% (w/v) SDS. Antibodies are listed in the Resource Table (Table 1). Membranes were imaged using an Odyssey® CLx Imaging System (LI-COR Biosciences) and band intensities were quantified using Image Studio™ Lite software (LI-COR Biosciences).

Band intensities were normalised to loading controls and presented relative to control conditions where indicated. Antibody specificity was validated by detection at the expected molecular weight and by loss of signal following RNA interference or genetic depletion, where applicable. Immunoblots shown are representative of several independent experiments.

### Co-immunoprecipitation

U2-OS cells stably expressing TCF25-FLAG or HEK-293T cells transiently expressing FLAG-tagged WT TCF25, ΔC^604-664^ TCF25 or empty pCMV-3Tag-3a backbone as a control were lysed in non-denaturing lysis buffer supplemented with protease inhibitors. Lysates were clarified by centrifugation, and protein concentrations were determined prior to immunoprecipitation.

Lysates were pre-cleared using a 1:1 mixture of Protein A/G magnetic beads (New England Biolabs) prior to enrichment. FLAG-tagged proteins were captured using anti-FLAG M2 magnetic beads (Sigma-Aldrich) under gentle rotation for 1 h at 4 °C. Following binding, beads were washed extensively in lysis buffer to remove non-specifically associated proteins. Immunoprecipitated proteins (10% of input) were denatured on beads in SDS sample buffer and analysed along with input by SDS–PAGE and immunoblotting.

### TurboID proximity labelling

The *TCF25* coding sequence was amplified from pCMV-3Tag-3a-TCF25 and cloned into the XhoI-BamHI-linearised pcDNA5/FRT/TO N-terminal 3×HA-TurboID expression vector using HiFi DNA assembly (New England Biolabs). Constructs expressing proximity-labelling fusion protein were propagated in *E. coli* and sequence-verified prior to use.

RPE-1 cells were transiently transfected with 3×HA-TurboID–TCF25, or the no-bait controls 3×HA-TurboID–∅ and 3×HA-TurboID–2×NLS, using FugeneHD. For proximity labelling, biotin-dependent labelling conditions were optimised, and 10 µM biotin for 1 h was selected to achieve robust enrichment of proximal biotinylated proteins. Cells were incubated with biotin, harvested by trypsinisation and collected in ice-cold PBS supplemented with protease inhibitor cocktail (Sigma-Aldrich, S8820), before snap-freezing cell pellet on dry ice. Pellet was lysed in modified RIPA buffer (50 mM Tris–HCl, pH 7.5, 150 mM NaCl, 1% NP-40, 0.5% SDC, 0.1% SDS, 1 mM EGTA, 1 mM EDTA) supplemented with inhibitors (as in WCL above) prior to streptavidin-based enrichment.

### Streptavidin enrichment and trypsin digestion

Biotinylated proteins were enriched using magnetic streptavidin beads (ReSyn Biosciences). Clarified lysates (0.4 mg protein) were incubated with equilibrated beads (25 µL slurry) in modified RIPA buffer for 1.5 h at 4 °C with gentle rotation. Beads were washed extensively using a series of high-salt, detergent and urea-containing buffers to remove non-specifically bound proteins.

Following the final wash, bound proteins were subjected to on-bead digestion in 50 mM TEAB buffer using MS-grade trypsin (0.5 µg) overnight at 37 °C. Dimethylated streptavidin beads were used to minimise streptavidin-derived peptide contamination while maintaining enrichment efficiency. Peptides were subsequently collected for mass spectrometric analysis.

### Mass spectrometry

LC–MS/MS was supported by the Proteomics Technology Platform, Francis Crick Institute. Peptides were analysed by LC–MS/MS using a timsTOF Pro2 mass spectrometer (Bruker) coupled to an Evosep One liquid chromatography system (Evosep). Peptides were loaded onto Evotips and separated using the 60 samples-per-day method on an Evosep performance column maintained at 40 °C. Data were acquired in DIA-PASEF mode over an m/z range of 100–1700 and an ion mobility range of 0.6–1.6 1/K₀, with ramp and accumulation times of 100 ms using sequential DIA windows spanning precursor mass and ion mobility space.

Raw data were processed using DIA-NN v1.8 against the human UniProt reference proteome with default settings [55,62]. RT-dependent normalisation was enabled, match-between-runs was disabled, and precursor mass accuracy was set to 15 ppm. Peptide and protein identifications were filtered using the default false discovery rate (FDR) control implemented in DIA-NN.

### Proteomics data analysis

Protein intensity values exported from DIA-NN were analysed using Perseus [55,57]. Intensities were log₂-transformed and missing values were imputed prior to statistical analysis. Proteomic measurements were derived from two independent biological replicates per condition. Differential enrichment between experimental groups was interpreted alongside fold-change magnitude of Log₂ fold-change > 2 and reproducibility across replicates.

Gene Ontology (GO) Cellular Component over-representation analysis was performed using g:Profiler on the list of reproducible proximal TurboID–TCF25 candidates (Log2FC > 2 in both replicates). Multiple testing correction was performed using the Benjamini–Hochberg method. Significant localisation terms (FDR < 0.05) with at least three associated proteins were retained for visualisation. To improve interpretability, redundant parent terms and non-localisation terms were removed after enrichment.

### Computational modelling of protein abundance

Protein-abundance quantitative mass-spectrometry measurements were obtained from the DepMap Harmonized Public Proteomics 24Q4 release of the CCLE proteomics study [36]. Protein identifiers were harmonised to UniProt primary accession numbers and mapped to HGNC gene symbols and Entrez Gene identifiers. Cell-line metadata were obtained from DepMap Public 26Q1 release. For Bayesian regression analysis, cell lines with complete normalised TCF25 and GPRASP2 abundance measurements were included. Bayesian linear regression was implemented in PyMC v5.23.0 with Normal priors for the intercept and slope (mean 0; standard deviation 5) and a Half-Normal prior for the residual standard deviation (standard deviation 2). Posterior inference used NUTS with two Markov chains and 1,000 tuning iterations and 1,000 posterior draws per chain. Posterior predictive distributions were generated along the regression line and for each observed cell line. Posterior means and 5^th^-95^th^ percentile predictive intervals described model uncertainty, while posterior predictive z-scores were used for quantifying observation-level deviation from the fitted model.

For functional enrichment analysis, proteins were ranked by posterior predictive deviation from the Bayesian model and analysed using pre-ranked Gene Set Enrichment Analysis (GSEA) implemented in GSEApy v1.1.4. Enrichment was summarised using normalised enrichment scores.

### Protein-protein interaction networks

Protein-protein interaction (PPI) networks were visualised using STRING (v11.5), incorporating known and predicted protein–protein associations [21]. Candidate interactors identified from Predictomes, a resource of systematically predicted protein complexes, were incorporated into the network [25]. Where indicated, networks were restricted to experimentally supported physical interactions using STRING’s physical subnetwork filter, with a minimum confidence threshold of 0.4 (medium) [21]. BioGRID was used as a curated resource for identifying physical interactions [54].

### Structural modelling

Structural modelling of the TCF25–GPRASP2 complexes was performed using the AlphaFold 3 public server with standard parameters and Protenix-v2 [37,38]. Predicted models were evaluated using confidence metrics including pLDDT, pTM and ipTM scores and PAE plots. Structural visualisation and analysis were performed using UCSF Chimera [58]. Predicted interfaces were interpreted qualitatively and used to guide hypothesis generation.

### Statistical analysis

Statistical analyses were performed using GraphPad Prism (v9.4.1). Quantitative data are presented as mean ± standard deviation (SD), with the number of biological replicates and statistical tests specified in the corresponding figure legends. Where appropriate, data distribution was assessed for normality using the Shapiro–Wilk test. *p*-values ≤ 0.05 were considered statistically significant. Figures were assembled using Inkscape v1.3.2.

## Data availability

The TurboID proteomics data have been deposited to the MassIVE repository (MassIVE ID MSV000103297). The output files for the TCF25 peptide—GPRASP2 complex from both AlphaFold 3 and Protenix-v2 and additional data are available upon request from the corresponding author.

## Supporting information

This article contains Supporting information.

## Acknowledgments

We are grateful to former members of the Khoronenkova laboratory, Madalena Castro and Mona Furukawa, for generating the *TCF25* knockout HEK293-T and U2-OS TCF25-FLAG cell lines, respectively. We thank Amelia Collins for providing hiPSC-derived cardiomyocytes and Tania Auchynnikava (Proteomics Technology Platform, Francis Crick Institute) for performing LC–MS/MS acquisition and DIA-NN data processing. We thank Marc de la Roche (University of Cambridge) for general support with tissue culture and cell models.

## Author contributions

Liana J.E. Hardy: conceptualisation, methodology, validation, formal analysis, investigation, data curation, visualisation, writing - original draft & editing. Haziq K. Moinudeen: methodology, formal analysis, investigation, data curation, visualisation. Laura Grzegorzek: methodology, validation, investigation. Fernando J. Bazan: methodology, formal analysis, investigation, visualisation, writing - editing. Svetlana Khoronenkova: conceptualisation, validation, resources, data curation, visualisation, supervision, project administration, funding acquisition, writing - original draft, review & editing.

## Funding and additional information

Liana J.E. Hardy was supported by a CRUK Cambridge Centre PhD studentship. Research in the Khoronenkova laboratory was funded by a Wellcome Trust and Royal Society Sir Henry Dale Fellowship [107643/B/15/Z] and the Royal Society [RGS/R1/201043].

## Conflict of interest

The authors declare that they have no conflicts of interest with the contents of this article.

## Notes

### Competing Interest Statement

The authors have declared no competing interest.

### Summary of Updates

This version of the manuscript includes large-scale analyses of the correlation between TCF25 and GPRASP2 protein levels. It also presents improved structural predictions of the putative TCF25-GPRASP2 interaction interface and experimental validation of this interface using deletion mutagenesis.

