## Supplementary Material for "TCF25 emerges as a cytoplasmic regulator of GPRASP2 stability"

**Supplementary Figure S1** provides the analyses of TCF25 interaction networks based on publicly available data. **Supplementary Figure S2** describes the results of TurboID–TCF25 proximity labelling. **Supplementary Figure S3** provides structural modelling of the TCF25-GPRASP2 interaction supporting *Fig. 5A*. **Supplementary Table S1** provides the complete TurboID–TCF25 proximity labelling dataset and the list of reproducible proximal candidates supporting *Fig. S2*.

**Fig. S1. Visualisation of TCF25 interaction networks using publicly available data.**

*(A)* STRING network analysis of predicted TCF25-associated proteins. *Left:* complete network including functional and physical associations with confidence scores indicated [1]. *Right:* network of Predictome-supported candidate proteins, visualised using STRING and restricted to physical interactions using STRING's built-in "physical subnetwork" classification [1,2]. Only interaction evidence indicating membership of the same physical complex is included, with a combined confidence score ≥ 0.400 (medium confidence). Blue lines: reported evidence from curated experimental interaction databases (BioGRID, IntAct, MINT, DIP), including yeast two-hybrid and large-scale affinity purification–mass spectrometry datasets. Red lines: text-mining evidence restricted to literature statements describing direct physical binding or complex formation, not general co-occurrence. Dashed lines: Predictome SPOC score, a structural-interaction confidence score based on AlphaFold-Multimer [2]. TCF25 (pink) is the bait protein in both networks; GPRASP2 (orange) is the only candidate shared between the networks.

*(B)* Histogram showing the reproducibility of physical interactors of TCF25 across independent studies reported in the Biological General Repository for Interaction Datasets BioGRID [3].

Physical interactions reported in a single independent study, such as LRRK1, were filtered out.

**Fig. S2. Identification of proteins proximal to TCF25 using TurboID labelling.**

*(A)* Schematic of TurboID proximity labelling workflow, including biotin-dependent labelling, streptavidin enrichment and LC–MS/MS analysis. Controls: no ligase (transfection reagent only), TurboID–∅ (whole-cell no-bait control plasmid), TurboID–2×NLS (nuclear no-bait control plasmid). Biotinylation profiles of TurboID–∅ and nuclear no-bait TurboID–2×NLS controls were largely similar, while their comparison with the no-ligase control defined the background biotinylation landscape for filtering non-specific hits in downstream analyses.

*(B)* Volcano plot of proteins identified by LC–MS/MS following TurboID–TCF25 proximity labelling in RPE-1 cells as in *Table S1* (n = 2 independent experiments, A and B). TurboID–∅ samples were used to define the biotinylation background; TurboID–2×NLS served as a nuclear localisation control. As TCF25 is predominantly cytoplasmic, enrichment was assessed relative to TurboID–∅ controls. Log$_2$ fold change (TurboID–TCF25/TurboID-∅) of experiments A *vs* B shown, with proteins with Log$_2$FC > 2 in both experiments highlighted in top right quadrant as reproducible candidates. No statistical analysis included due to the limited number of replicates. TCF25 (bait), the top 10 proximal proteins and selected proteins previously implicated in TCF25 biology are annotated.

*(C)* Gene Ontology (GO) Cellular Component analysis of reproducibly enriched TurboID–TCF25 candidate proteins (Log$_2$FC > 2 in both biological replicates) as in *(B)*. Significant GO terms defined by false discovery rate (FDR < 0.05). For visualisation, redundant GO localisation terms were consolidated into representative higher-level localisation categories while preserving proteins annotated to multiple cellular compartments.

**Fig. S3. Modelling the interaction of full-length TCF25 and GPRASP2 with refinement to predicted globular domains for further analysis.**

Predicted structure of the full-length TCF25-GPRASP2 complex with predicted template modelling (pTM) and interface predicted TM (ipTM) scores *(left)*. Predicted aligned error (PAE) plot *(top right)* and AlphaMissense variant-effect profiles *(bottom)* support the prediction of putative globular domains in TCF25 and GPRASP2 (dashed boxes). Predicted interaction surfaces are highlighted (red boxes). Predictions generated using the AlphaFold v3.2 Server [4].

**Table S1. TurboID–TCF25 proximity labelling proteomics dataset.**

Complete LC–MS/MS dataset from TurboID-mediated proximity labelling experiments comparing TurboID–TCF25 with control samples. The table includes raw intensities and $\text{Log}_2$ fold change (FC) values for all identified proteins. These proteins were used for volcano plot visualisation in *Fig. S1B*.
