## Supplementary figures and images for "TCF25 emerges as a cytoplasmic regulator of GPRASP2 stability"

### Supplementary Figures S1-S3

**A**

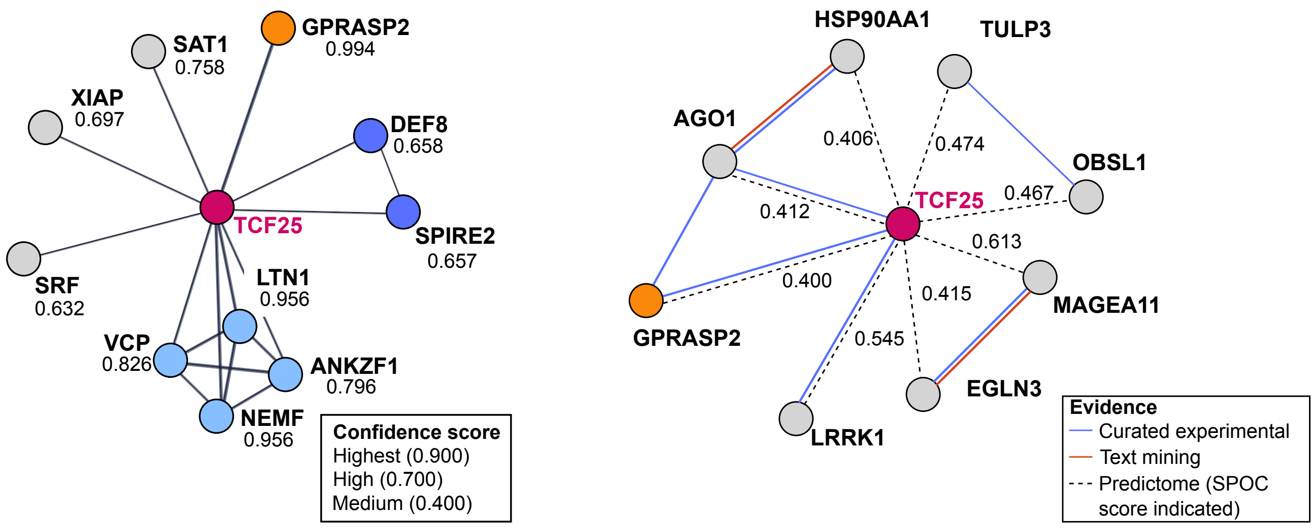


**B**

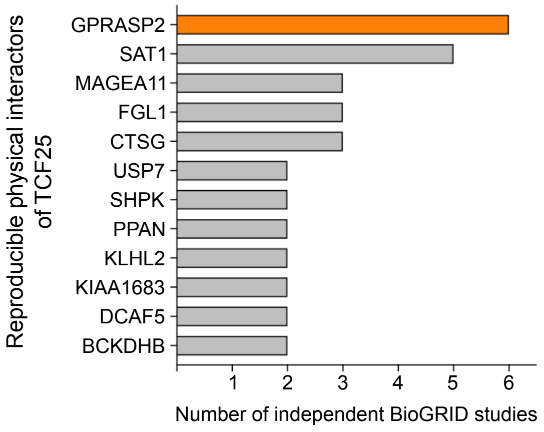


Hardy *et al.*, Figure S1

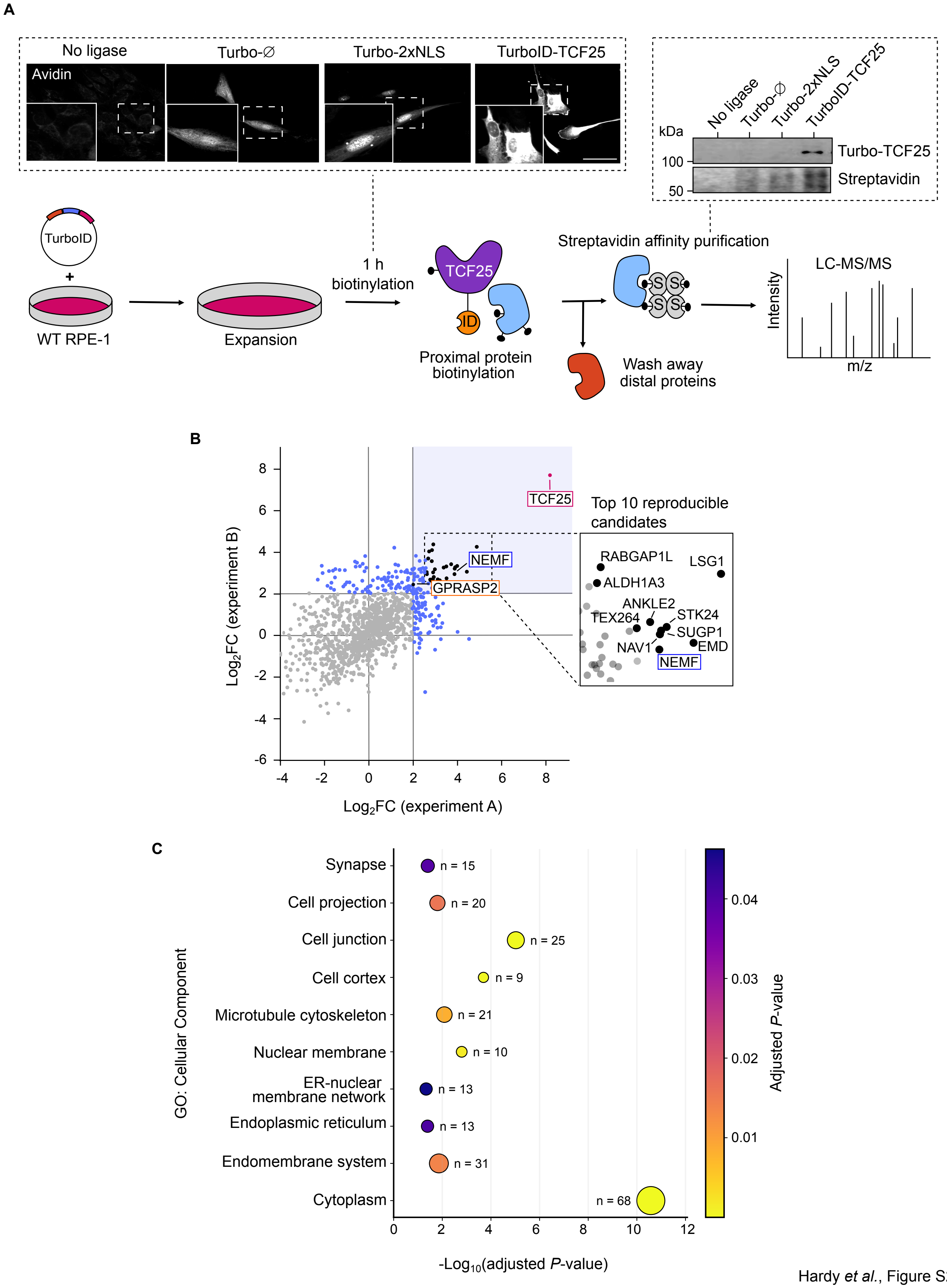


Hardy *et al.*, Figure S2

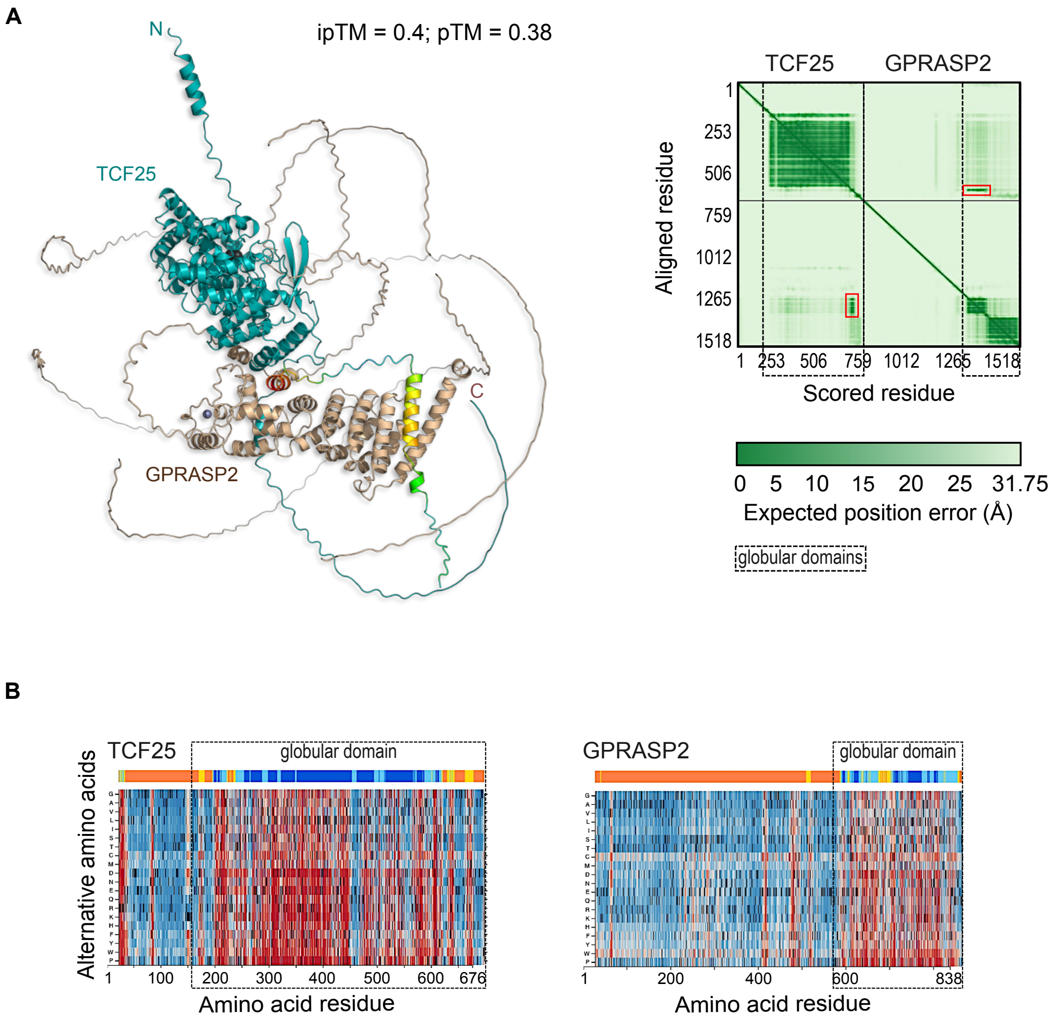


Hardy *et al.*, Figure S3
